# Electroporated recombinant proteins as tools for *in vivo* functional complementation, imaging, and chemical biology

**DOI:** 10.1101/631358

**Authors:** Amal Alex, Valentina Piano, Soumitra Polley, Marchel Stuiver, Stephanie Voss, Giuseppe Ciossani, Katharina Overlack, Beate Voss, Sabine Wohlgemuth, Arsen Petrovic, Yao-Wen Wu, Philipp Selenko, Andrea Musacchio, Stefano Maffini

## Abstract

Delivery of native or chemically modified recombinant proteins into mammalian cells shows promise for functional investigations and various technological applications, but concerns that sub-cellular localization and functional integrity of delivered proteins may be affected remain high. Here, we surveyed batch electroporation as a delivery tool for single polypeptides and multi-subunit protein assemblies of kinetochores, a spatially confined and well-studied subcellular structures. After electroporation in human cells, recombinant fluorescent Ndc80 and Mis12 multi-subunit complexes displayed native localization, physically interacted with endogenous binding partners, and functionally complemented depleted endogenous counterparts to promote mitotic checkpoint signaling and chromosome segregation. Farnesylation is required for kinetochore localization of the Dynein adaptor Spindly. In cells with chronically inhibited farnesyl transferase activity, *in vitro* farnesylation and electroporation reconstituted robust kinetochore localization of Spindly. Thus, electroporation is uniquely versatile for delivering synthetic and, as required, chemically modified functional mimics of endogenous proteins, and is therefore a promising tool for chemical and synthetic biology.

## Introduction

In molecular cell biology, the traditional separation between *in vivo* (or at least *in cellulo*) and *in vitro* approaches has been progressively surpassed in recent years thanks to new conceptual and technical advances. For instance, dissection and biochemical reconstitution of complex cellular pathways *in vitro* demonstrated great predictive value, facilitating and directing functional characterizations aiming to understand the endogenous cellular counterpart (e.g. (1, 2)). Chemical and synthetic biology, on the other hand, have generated tools for studying and manipulating biological pathways in creative new ways, with innovations that include, amongst others, genetic code expansion to introduce unnatural aminoacids with new functionalities into a protein of interest (3), the development of fluorescent dyes with potential for single molecule sensitivity and spanning a greater spectral range compared to genetically encoded proteins (4), and the design of allelic mutants of crucial enzymes that are specifically inhibited by small-molecules that won’t target the wild type allele (5).

Enclosure of the cell’s inner structures within biological membranes hinders the cellular delivery of soluble (macro)molecules and challenges the alliance of chemical and synthetic biology with molecular cell biology. Accurate delivery of surgical probes inside cells with the aim of dissecting complex intra-cellular pathways remains an art. Historically, introduction of protein variants into cells has been achieved by transient transfection of DNA plasmids encoding these entities. Recipient cells then typically express the desired proteins at levels that vary between cell lines and individual cells (6). Owing to the ease of transient transfection procedures, the method is widely used. Stable transfection of cultured mammalian cells offers more uniform protein expression but the generation of stable clones is labor-intensive and time-consuming (6).

A significant fraction of cellular proteins exists within stable or dynamic multi-subunit assemblies (7–9). Functional characterization of protein complexes in mammalian cells remains laborious especially when multiple subunits need to be manipulated (e.g. mutations or the introduction of fluorescent probes) for functional complementation studies. In such cases, multiple subunits must be co-expressed to similar levels while endogenous counterparts ought to be depleted, which is technically challenging and exposes a potential limitation of direct DNA-based delivery methods. Because tools for recombinant expression of protein complexes in heterologous systems, including bacteria and insect cells, are abundantly available in structural biology (10), the prospects of directly delivering these specimens into cells are highly attractive. In addition, such procedures enable the establishment of defined protein modifications before intracellular delivery, which allows researchers to exploit the repertoire of chemical biology tools for in-cell investigations.

The tools available for protein delivery, however, remain insufficiently explored (11). Historically, microinjection first emerged as the method of choice for the introduction of recombinant proteins into mammalian cells (12). However, it requires considerable handling skills and a dedicated setup that are not routinely available, and is a low-throughput approach that limits delivery to a small number of targeted cells (typically in the range of tens to hundreds). To achieve higher throughput, batch delivery methods such as cell-penetrating peptide (CPP) tags (13), pore-forming bacterial toxins (14, 15) or osmocytosis (16) are available, but present uncertainties with regard to protein uptake efficiencies, undefined routes of internalization and intracellular localization, as well as varying degrees of cytotoxicity.

Batch electroporation (EP) is an alternative method to directly deliver proteins into cultured mammalian cells (17–20). EP-mediated protein transduction has surged in recent years with applications ranging from, cellular structural biology and the delivery of isotopically labelled proteins for in-cell NMR measurements (21) to the introduction of immunoglobulins for targeting and degradation of endogenous proteins (22), and the delivery of ribonucleotide particles (RNPs) for CRISPR-Cas9-mediated gene editing (23–26).

While these approaches provide a promising outlook for future applications, more experimental evidence regarding the structural and functional intactness of delivered proteins in cells and a comprehensive analysis of the robustness and reproducibility with which different mammalian cells can be targeted is needed. Here, we test EP as a delivery method for various proteins that work in the kinetochore, a multi-subunit assembly required for chromosome segregation (27). The kinetochore consists of at least 30 core subunits organized in various protein assemblies and structured in layers that emanate from a specialized centromeric chromatin interface, the inner kinetochore, and terminate at the outer kinetochore, where spindle microtubules are bound (Figure 1A). The primary function of kinetochores is to link chromosomes and spindle microtubules (MTs) (28). Kinetochores function also as signaling platforms for the spindle assembly checkpoint (SAC, also named mitotic or metaphase checkpoint), a feedback surveillance mechanism that monitors the correct state of kinetochore-microtubule (KT-MT) attachment to ensure timely and faithful chromosome segregation (29).

**Figure 1.**
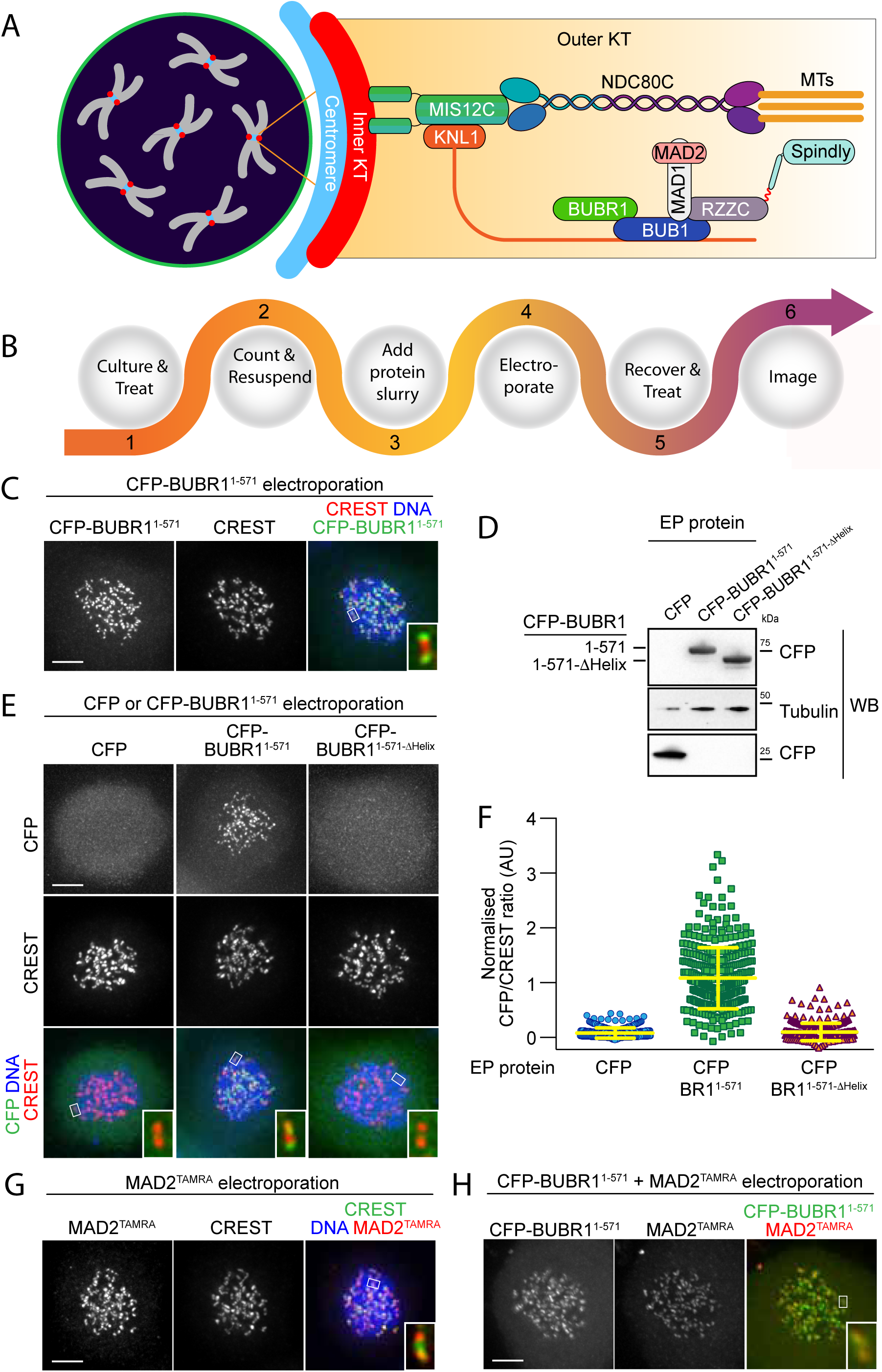
Delivery of spindle assembly checkpoint proteins by electroporation. **A**) Schematic outline of a mitotic cell with condensed chromosomes (grey) and centromeres (blue). The inner kinetochore (red) recruits several outer kinetochore proteins (orange gradient box), including the MIS12 (MIS12C) and the microtubule (MT) binding complex NDC80 (NDC80C). In turn, these complexes recruit the protein KNL1, the spindle assembly checkpoint (SAC) components BUB1, BUBR1, MAD1 and MAD2, and the outer kinetochore proteins RZZ and Spindly. **B**) Overview of the electroporation (EP) work-flow using adherent cells. (1) Cells are cultured under defined growth conditions according to experimental requirements. (2) Prior to EP, cells are harvested by trypsinization, washed in PBS, counted and resuspended in EP buffer. (3) Recombinant protein diluted in EP buffer is added to cell slurries at desired concentrations (this stage is called EP slurry). (4) Cells are pulsed between 1 and 3-times to allow for efficient delivery of recombinant proteins. (5) Cells are tripsinised and then washed twice to remove non-incorporated protein, and then plated on cover slips or culture flasks to allow for recovery. (6) After a recovery period, cells are processed for imaging. **C**) Following EP of CFP-BUBR1^1-571^ and over-night recovery, cells were treated for 6 hours with nocodazole and then prepared for immunofluorescence analysis. Kinetochores were stained with the marker CREST and DNA with SiR-Hoechst-647. Insets represents magnifications of the boxed kinetochores. Scale bar = 5 µm. **D**) CFP alone, CFP-BUBR1^1-571^ and the BUB1-binding-deficient mutant CFP-BUBR1^1-571-ΔHelix^ were electroporated under the same conditions used in C. Protein extracts generated from electroporated cells were subjected to western blotting analysis with the indicated antibodies. **E**) CFP-BUBR1^1-571-ΔHelix^ fails to localize to kinetochores. Kinetochores were stained with the marker CREST and DNA with SiR-Hoechst-647. Scale bar = 5 μm. **F**) Quantification of KT levels for electroporated CFP proteins from cells shown in E. Each symbol represents a single cell. Yellow lines indicate median values ± interquartile range. **G)** SAC protein MAD2^TAMRA^ electroporated into cells display proper localization to unattached kinetochores. Kinetochores were stained with the marker CREST and DNA with DAPI. Scale bar = 5 μm. Insets represent magnifications of the indicated KT. **H)** Simultaneous electroporation of CFP-BUBR1^1-571^ and MAD2^TAMRA^ results in proper localization to kinetochores of nocodazole treated HeLa cells. Insets represent magnifications of the indicated kinetochore. Scale bar = 5 µm.

The advantage of using kinetochores in this study is that they form small, discrete cellular structures with a very recognizable punctuate pattern and several markers are available to probe their native composition, thus making it very easy to assess correct localization of exogenously introduced variants. Therefore, correct assembly of kinetochore complexes upon subunit delivery by EP can be inspected at different levels of structural and functional resolution (27). Our results demonstrate that electroporation is well suited to deliver individual kinetochore proteins into different mammalian cell types and that they assemble properly into native kinetochore complexes. We further show that electroporation can be used to transduce pre-assembled kinetochore components to functionally complement depleted KT subunits. Collectively, our results provide very encouraging evidence that EP can be used to deliver recombinant proteins of interest, even when chemically modified, into target cells for functional complementation in a non-invasive manner.

## Results and Discussion

### Electroporation: an outline

As outlined in Figure 1B and as described in detail in the Methods section, intracellular protein delivery by EP is a simple procedure that does not require specialized handling skills or dedicated instrumental set-ups, making it easy to establish in any laboratory. Briefly, in preparation for EP, cells can be treated under various experimental regimes, e.g. for cell cycle synchronization; (30) before being harvested, washed, and resuspended in an electroporation buffer containing the diluted protein (the “EP slurry”). The EP slurry is subjected to multiple electric pulses that create reversible ruptures in the cell’s membrane and allow intracellular protein delivery. Following EP, cells are treated with trypsin and washed to remove protein aggregates from the outer side of the cell membrane and allowed to recover, typically between 16 and 18 hours but earlier times are possible (e.g. 4 hours), before being prepared for the desired analysis, such as live imaging, immunofluorescence, immunoprecipitation or Western blotting. Using model recombinant protein like α-Synuclein (made fluorescent by labeling with the synthetic dye Atto488) or mCherry, which can reach very high concentrations, we observed linear uptake characteristics over a broad range of input protein concentrations in the EP slurry (Figure S1A-B), indicating that EP allows for semi-quantitative delivery of recombinant proteins. In turn, this allows for the generation of cell populations harboring defined amounts of exogenously delivered proteins (Figure S1C-D). Importantly, EP is efficient across multiple cell lines and treated cells show little or no sign of damage or toxicity (Figure S1A and D-F).

Previously, we generated various recombinant versions of stable sub-complexes of the human kinetochore, and demonstrated that they could be used to reconstitute large parts of the kinetochore and of the spindle assembly checkpoint (SAC) *in vitro* (1, 28, 31, 32). This library of high-quality recombinant samples (Figure S2) already extensively characterized *in vitro* represented an excellent resource to address the general suitability of EP for *in vivo* delivery and functional analysis.

We initially tested this approach on BUBR1 (BUB1-related), a multi-domain protein that operates in the SAC. Recombinant BUBR1^1-571^ (encompassing residues 1-571 of the 1050-residue BubR1 protein, which are sufficient for checkpoint function) (1) fused to a cyan fluorescent protein (CFP) was introduced into HeLa cells at an EP slurry concentration of 10 μM. In mitotically arrested cells (by addition of the spindle poison nocodazole, a condition that activates the SAC), CFP-BUBR1^1-571^ distinctly localized to kinetochores, as revealed by the typical punctuate staining and by co-localization with CREST, an established kinetochore marker (Figure 1C). We have shown previously that BUBR1 kinetochore recruitment in human cells requires dimerization with kinetochore-bound BUB1, a BUBR1 paralog, through segments that encompass a predicted helical domain in the two proteins (33). In agreement with these previous observations, recombinant CFP-BUBR1^1-571-ΔHelix^, a deletion mutant of BUBR1 lacking the BUB1 binding helix (deletion of residues 432-484) was unable to localize to kinetochores (Figure 1D-F). Collectively, these results demonstrate that, following EP, recombinant CFP-BUBR1^1-571^ binds to its endogenous kinetochore receptor BUB1 in target cells. The efficiency of CFP-BUBR1^1-571^ delivery was high and most cells displayed uniform levels of the recombinant protein (Figure S3). Thus, electroporation can deliver very similar amounts of recombinant protein into each cell. In the course of this study, this became to be recognized as a general feature of this approach, largely independently of the protein used.

We used the Sortase method (34) to fuse a fluorescent peptide modified with the dye tetra-methyl-rhodamine (TAMRA) to the C-terminus of MAD2, another SAC protein. The resulting MAD2^TAMRA^ was delivered by EP in HeLa cells at an EP-slurry concentration of 5 μM. Like CFP-BUBR1^1-571^, MAD2^TAMRA^ was found to localize effectively at kinetochores of nocodazole-treated cells, as shown by co-localization with CREST (Figure 1G). Furthermore, we performed co-delivery of CFP-BUBR1^1-571^ with MAD2^TAMRA^, and found both proteins at unattached KTs in nocodazole-arrested cells (Figure 1H). Thus, individual or multiple fluorophore-tagged components of the human KT are faithfully targeted to their respective binding sites after EP.

In addition to localization studies, we tested if EP can be used in combination with in-cell spectroscopic applications, such as FLIM-FRET microscopy (35), to study the spatiotemporal dynamics of recombinant proteins *in vivo*. KRAS/p21 is a small, globular GTPase with important regulatory functions in many signaling pathways (36). We studied KRAS signaling in live cells electroporated with GFP-tagged and Tide Fluor 3 (TF3)-coupled human KRAS (i.e. EGFP-TF3-KRAS, final size ~50 kDa), a conformational-change FRET sensor purified from *E. coli* (37) (Figure S4A). Activation and conformational switching of EGFP-TF3-KRAS occurs upon epidermal growth factor (EGF) stimulation and causes the spatial separation of the fluorescence moieties and a concomitant loss of intramolecular FRET signal, which ultimately results in an increase of the donor fluorescence lifetime measured by FLIM-FRET live microscopy (37). Upon EP-delivery, we found that EGFP-TF3-KRAS, but not EGFP, predominantly localize to the plasma membrane (Figure S4B), with a pattern indistinguishable from endogenous, over-expressed, or microinjected KRAS (36–38). Following addition of EGF, we observed the predicted increase in the donor lifetime of EGFP-TF3-KRAS (Figure S4C-D). Importantly, our EP-based results were qualitatively and quantitatively identical to the ones that were observed in EGFP-TF3-KRAS microinjection experiments (37), demonstrating the suitability of EP for FRET-based assays in living cells as a valid alternative to protein delivery by microinjection.

### Electroporated MIS12 complex targets kinetochores and functionally complements depletion of the endogenous complex

The experiments discussed in the previous section described delivery of single polypeptides. Next, we tested the suitability of EP for delivery of multi-subunit protein complexes. To this end, we started with the Mis12 complex (MIS12C, where ‘C’ stands for ‘complex’), an extremely stable four-subunit kinetochore assembly of the DSN1, NSL1, MIS12, and PMF1 subunits with molecular mass of 120 kDa. In previous studies, we demonstrated that the stability of the MIS12C requires co-expression of all four constituent subunits and reported the development of various co-expression vectors for its heterologous reconstitution in insect cells (39–41). Here we generated a fluorescence version of MIS12C by fusing the coding sequence of cyan fluorescent protein (CFP) to that of the PMF1 subunit. We then co-expressed the subunits in insect cells and purified the resulting complex (CFP-MIS12C) to homogeneity (Figure S2).

We electroporated CFP-MIS12C into cycling HeLa cells and monitored CFP-MIS12C levels by fluorescent microscopy. At an EP slurry concentration of 8 μM, CFP-MIS12C was clearly visible at kinetochores in the large majority of electroporated cells and with remarkably similar kinetochore levels within the same cell and between different cells (Figure 2B and Figure S3C). The ability of the electroporated CFP-MIS12C to reach kinetochores after introduction in the cellular environment indicates that the complex retains its integrity through the electroporation process.

**Figure 2.**
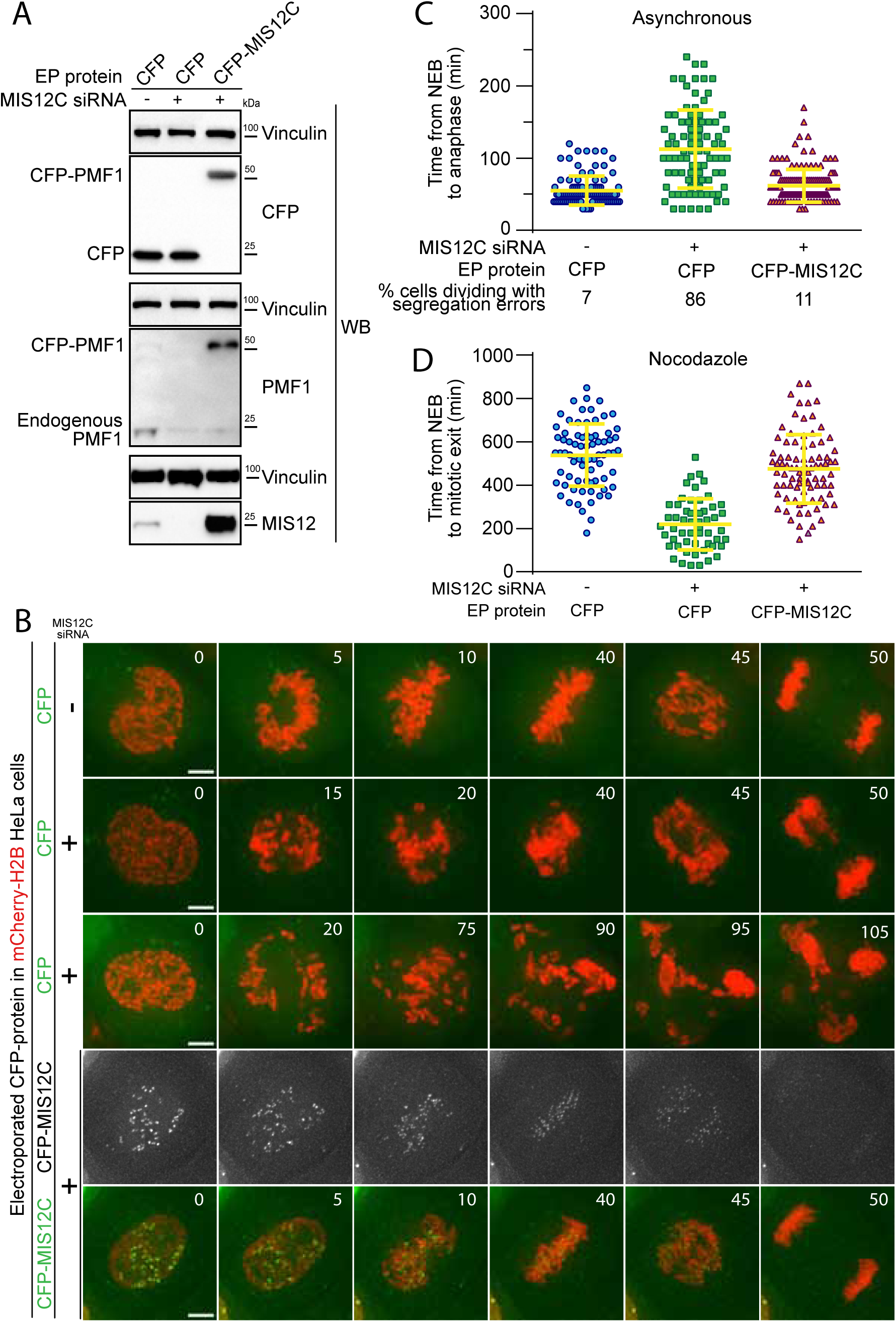
Electroporated MIS12 complex targets kinetochores and functionally complements depletion of the endogenous complex. **A**) The indicated proteins, including CFP-labeled recombinant MIS12C, were electroporated in mCherry-H2B-expressing HeLa cells previously treated with siRNAs to deplete the endogenous MIS12C. Western blotting of cellular protein levels in extracts was performed and showed a depletion in MIS12 and PMF1 levels of 93,7% and 87,6% respectively. **B**) Frames collected at the indicated time points (min) of time-lapse live-cell fluorescence microscopy movies depicts chromosome congression and segregation of cells from A. Two representative phenotypes observed in cells with depleted MIS12C that were concomitantly electroporated with CFP as control are shown. Green and red signal represents, respectively, CFP proteins and DNA. Scale bar = 5 µm. **C**) Quantification of chromosome congression and segregation defects in B. Each symbol represents a single cell. Yellow lines indicate mean values ± SD. **D**) Electroporated CFP-MIS12C rescues SAC defects in H2B-mCherry HeLa cells depleted for endogenous MIS12C. Duration of mitosis (based on DNA and cell morphology) of individual cells treated as in A but imaged in the presence of nocodazole 0.3 μM was plotted. Yellow lines indicate mean values ± SD.

In a more stringent test, we also asked if CFP-MIS12C retained its expected kinetochore function after electroporation. MIS12C is as an assembly hub for the kinetochore, being responsible for the recruitment of a subset of the microtubule-binding NDC80 complexes as well as of KNL1, a signaling platform for the SAC (41, 42) (Figure 1A). These functions make MIS12C necessary for timely chromosome congression and faithful cell division. To test the functionality of the recombinant CFP-MIS12 complex, we performed a complementation assay in HeLa cells depleted of endogenous MIS12 complex by the RNA interference (RNAi) methods using silencing RNAs (siRNAs) against multiple subunits, as previously described (43). Effective depletion of MIS12C subunits and uptake of fluorescent proteins was demonstrated by Western blotting (Figure 2A). This is noteworthy, as it exposes a fundamental advantage of EP over another protein targeting technique such as microinjection, which is limited to a small number of target cells and therefore unsuitable to biochemical analyses. In agreement with previous reports, depletion of MIS12C compromised chromosome congression (Figure 2B-C), a consequence of reduced kinetochore levels of NDC80 and KNL1 (43). As shown by live-cell microscopy, EP delivery of CFP-MIS12C into depleted cells, but not of a CFP control, rescued chromosome congression essentially to the levels observed in control cells, indicating that the recombinant complex is functionally active (Figure 2B-C and **Supplementary movies** 1-4).

Improperly attached kinetochores trigger the spindle assembly checkpoint (SAC), causing a mitotic arrest that delays entry into anaphase (29). The checkpoint pathway exhausts its function only after all chromosomes have achieved bi-orientation on the mitotic spindle. Despite causing dramatic chromosome congression failures, depletion of endogenous MIS12C is hardly compatible with checkpoint function because cells lacking the MIS12C also fail to recruit KNL1, which is required for SAC signaling. Thus, MIS12C depleted cells have a strongly weakened SAC that results in their premature anaphase and mitotic exit (43, 44). To assess SAC function under our RNAi conditions, we treated cells with 0.3 μM of the microtubule-depolymerizer nocodazole, thus creating conditions that potently activate checkpoint signaling in control cells. We then measured, by live-cell video microscopy, the cells’ ability to sustain a prolonged mitotic arrest. In agreement with the previous studies, cells depleted of MIS12C were unable to mount a strong mitotic arrest and underwent mitotic exit in presence of multiple unattached or incompletely attached chromosomes, indicative of a checkpoint defect. Electroporated CFP was unable to restore the SAC response, while recombinant CFP-MIS12C restored mitotic duration to values similar to those observed in control cells, indicating full rescue of SAC activity (Figure 2D). Collectively, these results indicate that electroporated recombinant MIS12C targeted the KT as a functionally intact complex and that EP can be used for biological complementation assays in combination with knockdown or knockouts.

### EP of a protein complex with an exceptionally large hydrodynamic radius and its interaction with endogenous partners

With a long axis of approximately 20 nm (39, 40) the MIS12C is rather elongated, a feature that at least qualitatively does not appear to interfere with efficient EP. With its dumbbell shape consisting of a coiled-coil shaft flanked by terminal globular domains, the four-subunit NDC80C complex reaches approximately 60 nm in length (31, 45). We were curious to assess if extreme elongation of NDC80C affected EP efficiency. As for the MIS12C, also the ~180 kDa NDC80C achieves stability through co-expression of its four subunits (NUF2, SPC24, SPC25, and NDC80) in insect cells ((40, 45); Figure 1A). After expression and purification with an established protocol of a fluorescent version of the complex containing NDC80-GFP (Figure S2), we electroporated it into cycling HeLa cells at an EP slurry concentration of 5 µM. 12 hours after electroporation, cells were imaged by live cell video-microscopy. This revealed kinetochore localization of the recombinant NDC80-GFP complex (Figure 3A), indicating that the considerable length of the complex is compatible with efficient delivery.

**Figure 3.**
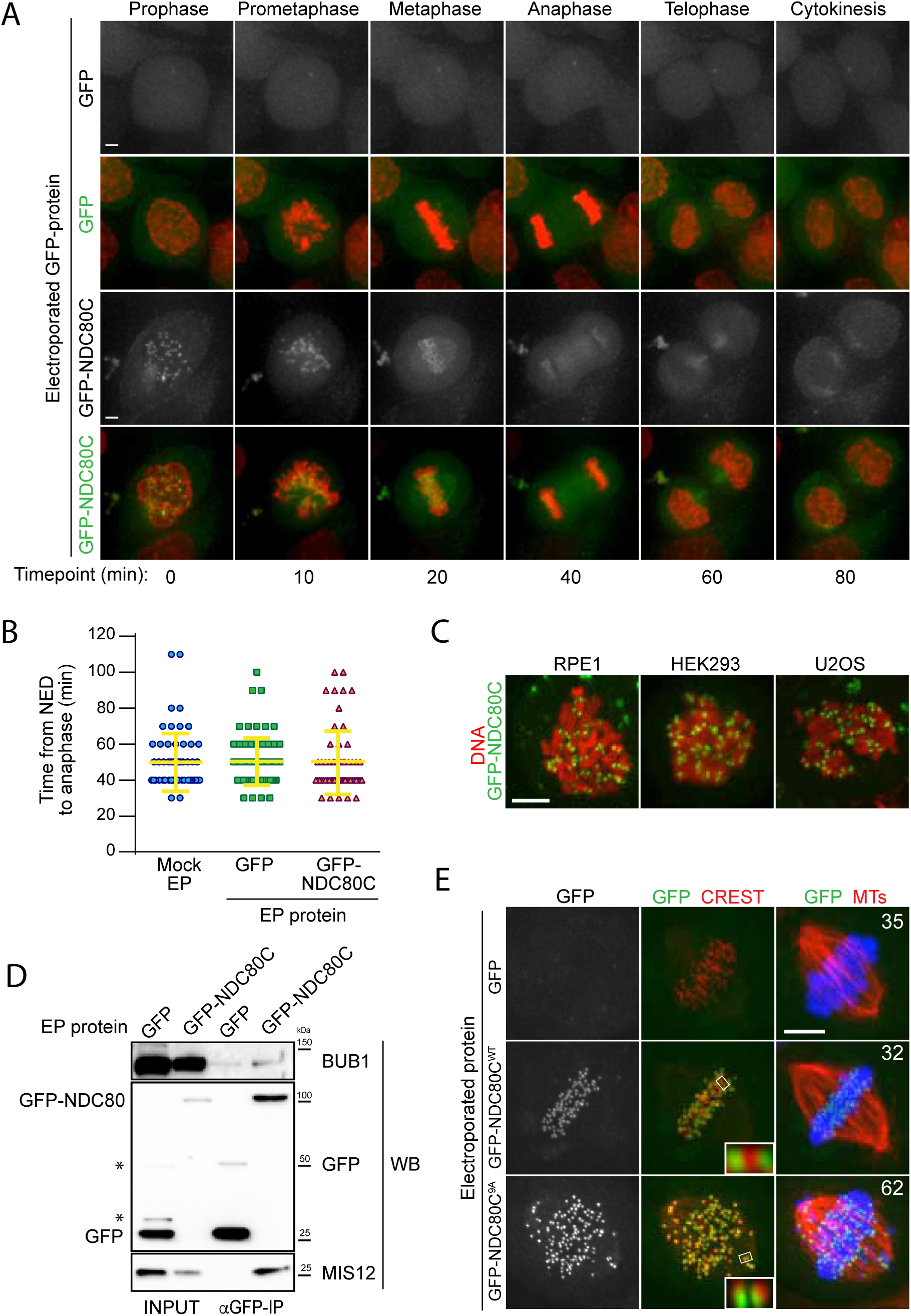
Cellular delivery of a protein complex with an exceptionally large hydrodynamic radius. **A**) GFP-labeled recombinant NDC80C^WT^ binds to kinetochores and its localization is compatible with normal mitotic progression. Panels show timepoints of time-lapse movies for asynchronously growing HeLa cells electroporated with recombinant GFP proteins and imaged live every 10 minutes. DNA was stained with SiR-Hoechst-647 and is shown in red. Scale bar = 5 µm. **B**) Quantification of time periods until anaphase onset for each of the conditions in panel A. Each symbol represents a single cell. Yellow lines indicate mean values ± SD. **C**) Additional human cell lines were electroporated with NDC80C-GFP. RPE1 cells are non-transformed human retinal pigmented epithelium cells. HEK293 cells are human epithelium kidney cells. U2OS cells are human osteosarcoma cells. Following recovery and staining with SiR-Hoechst-647 DNA dye, cells were imaged live and display a clearly visible kinetochore localization of the delivered NDC80C-GFP complex. Scale bar = 5 µm. **D**) Immunoprecipitation analysis of protein extracts from cells treated as in A shows that EP-delivered NDC80C-GFP^WT^ establishes normal interactions with its endogenous KT partners. Western blotting panels show immunoprecipitates performed with α-GFP beads and probed with the indicated antibodies. **E**) An error correction assay for the EP-delivery of a NDC80C^9A^-GFP mutant shows a dominant-negative effect on the correction of improper KT-MT attachments. Panels display representative images of electroporated HeLa cells treated with STLC to accumulate erroneous synthelic KT-MT attachments, followed by STLC washout, release into MG132 (proteasome inhibitor that prevents mitotic exit), and fixation 150 minutes after STLC-release. Inability to correct erroneous attachments results in uncongressed chromosomes. Control cells were electroporated with GFP alone. Numbers represent the percentage of cells with uncongressed chromosomes for each condition. KTs were labeled with anti-CREST immunostaining, microtubules (MT) with an anti-TUBULIN antibody and DNA with DAPI. Insets represent magnificatis of the indicated KT. Scale bar = 5 µm.

HeLa cells carrying recombinant NDC80C-GFP in addition to endogenous NDC80C displayed no alterations in cell cycle progression and proceeded through the different stages of mitosis without detectable changes when compared to control cells (Figure 3A-B and **Supplementary movies S**5-**S**9). Thus, delivering exogenous NDC80C-GFP does not cause obvious cell cycle alterations or cytotoxicity. When NDC80C-GFP was electroporated into other human cell lines, such as RPE, HEK293 and U2OS, equally clear kinetochore signals were observed (Figure 3C).

To test weather recombinant NDC80C-GFP complexes establish their expected physical interactions after EP delivery, we performed GFP-immunoprecipitation (GFP-IP) assays on protein extracts generated from mitotic HeLa cells previously electroporated with NDC80C-GFP. Western blotting demonstrated that MIS12 and BUB1 were present in the NDC80C-GFP precipitates, but not in precipitates of GFP (Figure 3D). Thus, the electroporated NDC80-GFP establishes physiologic interactions with its endogenous binding partners, demonstrating the versatility of EP as a technique to probe protein interactions in living cells both through imaging and through biochemistry.

### An NDC80C mutant triggers a dominant-negative chromosome alignment defect

Chromosome bi-orientation implies that the sister kinetochores attach to microtubules emanating from opposite spindle poles. Incorrect attachments that occur during mitosis are sensed by an error correction mechanism that, by regulating the phosphorylation state of the unstructured N-terminal region of the NDC80 subunit, causes the destabilization of incorrect attachment and the formation of new, correct ones (46). A previously described NDC80 mutant (9A) carrying nine alanine mutations at the phosphorylation sites in the protein’s N-terminus establishes hyper-stable KT-MT attachments that cannot be corrected, resulting in frequent chromosome congression errors (47–49). We asked if a NDC80C^9A^-GFP mutant electroporated in HeLa cells had dominant-negative effects in an established biorientation and error correction assay (50). HeLa cells were allowed to enter mitosis in presence of STLC, an inhibitor of the Eg5 kinesin (51) whose activity is crucially required for centrosome separation and spindle bipolarization, resulting in the accumulation of monopolar spindles in which sister kinetochores frequently attach to the same microtubule organizing center (MTOC, the centrosome in this case) in a condition known as synthelic attachment. Upon washout of STLC, Eg5 reactivation, and spindle bipolarization, a functional error correction pathway is required to resolve the synthelic attachments and achieve correct metaphase alignment (50). In this setup, control cells electroporated with GFP or NDC80C^WT^-GFP corrected erroneous attachments and aligned their chromosomes properly at the metaphase plate (Figure 3E). In contrast, cells electroporated with the NDC80C^9A^-GFP mutant were impaired in their correction pathway, and accumulated misaligned chromosomes with high frequency. Thus, recombinant NDC80C^9A^-GFP electroporated in HeLa cells exercises a dominant-negative effect on the ability of endogenous NDC80C to promote chromosome alignment.

### In vitro farnesylation allows Spindly localization when farnesyl transferase is inhibited

Spindly is an adaptor protein that promotes the interaction of the minus-end-directed motor Dynein with its processivity factor Dynactin (52, 53). Spindly also interacts directly with the ROD-Zwilch-ZW10 (RZZ) complex, the main constituent of the so-called kinetochore corona, a crescent-shaped structure that assembles on kinetochores in early prometaphase to promote microtubule capture (54) (Figure 1A). Kinetochore localization of Spindly critically depends on its ability to interact with the RZZ complex (55, 56). The latter, in turn, requires the irreversible post-translational isoprenylation of Spindly with a farnesyl moiety on cysteine residue near the Spindly C-terminus (55–57). Farnesylation is carried out by the enzyme farnesyl-transferase (FT), which, in addition to Spindly, also targets several members of a family of Ras-like small GTP binding proteins, including Ras itself (58). Potent FT inhibitors have been described, and previous studies demonstrated that the localization of Spindly to kinetochores is prevented when FT is inhibited (55, 56).

Because farnesylation of Spindly has been reconstituted *in vitro* with purified components (57), we reasoned that we could exploit this reaction to promote kinetochore localization of recombinant Spindly in cells experiencing a long-term blockade of FT activity. Two versions of HsSpindly that were N-terminally fused to GFP or mCherry (molecular mass ~95 kDa) were expressed in insect cells and purified to homogeneity (Figure S2). Our previous mass spectrometry analysis demonstrated that only trace farnesylation of Spindly takes place in insect cells (57). When we EP-delivered fluorescent recombinant mCherry-Spindly into HeLa cells, we detected a fluorescent signal at the periphery of kinetochores that was comparable to the localization described for endogenous Spindly (Figure 4A-B) (55), or of Spindly introduced by microinjection (unpublished observation). mCherry-Spindly^C602A^, a mutant that cannot be farnesylated, failed to show any KT localization upon EP (Figure 4A-B). This indicated that wild type mCherry-Spindly becomes farnesylated after intracellular delivery. To further substantiate this, we treated cells with FTI-227, a selective farnesyl-transferase inhibitor (58). In presence of FTI-227, localization of recombinant mCherry-Spindly or GFP-Spindly was greatly reduced (Figure 4B-C), confirming that FT activity is required for Spindly localization.

**Figure 4.**
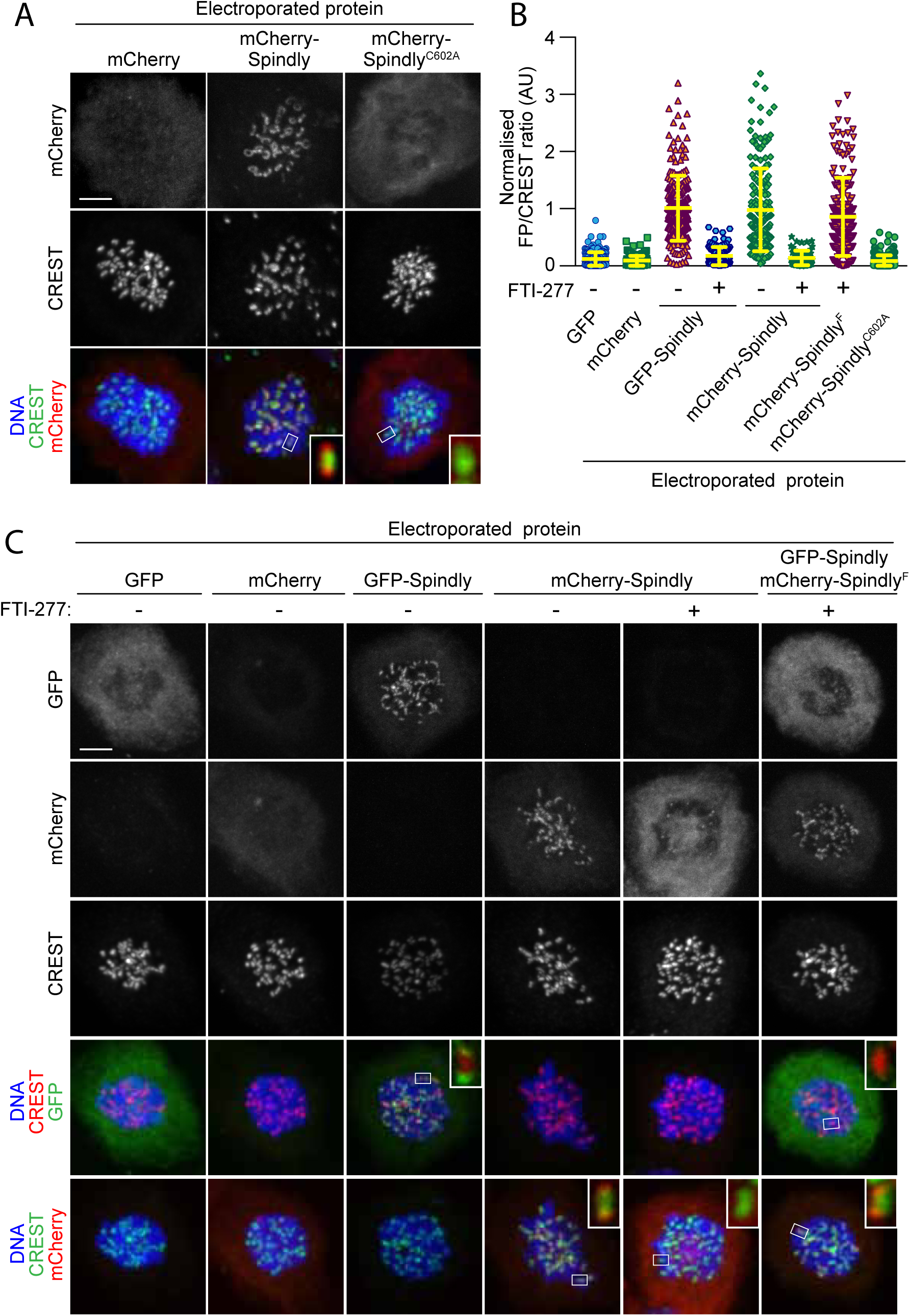
In vitro farnesylation allows Spindly localization when farnesyl transferase is inhibited. **A**) Kinetochore localization of unmodified recombinant Spindly depends on cellular farnesylation. Panels show representative images of cells transduced with unfarnesylated mCherry-Spindly in nocodazole-treated HeLa cells. Mutation of Cys602 to alanine prevents farnesylation upon EP and reduces KT levels of the delivered protein. Kinetochores were stained with CREST antibodies and DNA with DAPI. Controls cells were electroporated with mCherry. Insets represent magnifications of the indicated kinetochores. Scale bar = 5 µm. **B**) Quantification of KT levels for electroporated Spindly from cells in A and C. Each symbol represents a single cell. Yellow lines indicate mean values ± SD. **C**) I*n vitro* farnesylation of mCherry-Spindly before EP bypasses the need of cellular farnesylation to achieve kinetochore localization. Cells treated with the farnesyl-transferase inhibitor FTI-277 were electroporated with either GFP, GFP-Spindly, mCherry, mCherry-Spindly, or i*n vitro pre-*farnesylated mCherry-Spindly (mCherry-Spindly^F^) and then processed for immunofluorescence analysis. Kinetochores were stained for CREST and DNA with DAPI. Insets represent magnifications of the indicated KT. FP= Fluorescent protein. Scale bar = 5 µm.

We therefore asked if *in vitro* farnesylation by FT rescued the localization of Spindly in cells treated with FTI-227. mCherry-Spindly was farnesylated *in vitro* with recombinant farnesyltransferase (FT) and farnesyl-pyrophosphate as a substrate, as described previously (57). The resulting farnesylated mCherry-Spindly (mCherry-Spindly^F^) and untreated GFP-Spindly were co-electroporated at the same EP slurry concentration (10 µM) in HeLa cells that had been treated with FTI-227 for the previous 24 hours. Remarkably, mCherry-Spindly^F^, but not GFP-Spindly, successfully targeted kinetochores, showing that the complementation of FT activity *in vitro* had bypassed the blockade of FT activity in cells by FTI-227. Although we do not provide formal proof, the persistent treatment with FT inhibitors in this experiment suggests that Spindly^F^ may be the only residual pervasively farnesylated protein in the target cells.

## Conclusions

The modification of biological macromolecules through the introduction of new functionalities has progressed at a tremendous pace in recent years, enriching the palette of tools for biological investigation with a wide spectrum of functionalities. For instance, genetic code expansion, an approach that allows to introduce old and new chemical functionalities into proteins, from synthetic, photostable fluorescent dyes with high quantum yields to highly reactive activatable groups (e.g. UV-activated cross-linkers and click chemistry tools), is undergoing an explosive development (3, 59). With rapid growth of chemical and synthetic biology, the demand for robust methods to deliver modified synthetic or semi-synthetic macromolecules into cells is growing (60).

While electroporation has been long recognized for its potential as a delivery approach, questions have been raised regarding the degree of its invasiveness and damage inferred on cellular structures, most notably membranes (61). Here, we have assessed the suitability of batch EP for delivering recombinant proteins into living mammalian cells for various applications in cell biology. In comparison to previous studies that focused on cytosolic proteins devoid of specific localization, our focus was on kinetochores, which are small, discrete subcellular structures. This was crucial, because it enabled a detailed assessment of the effective degree of functional complementation of the electroporated samples. Our results identify EP as a rapid, efficient, and semi-quantitative technique that enables the delivery into various cultured mammalian cell lines of proteins of variable mass and hydrodynamic radius, including the highly elongated NDC80C. Importantly, we provide clear evidence that the delivered proteins remain structurally and functionally intact after delivery, as witnessed by their correct kinetochore localization and, where applicable, ability to complement RNAi-based depletion, dominant negative effects, and immunoprecipitation with endogenous interacting partners. Thus, in addition to its ease of application, EP is very attractive because it allows delivery in sufficiently large cohorts of cells, and is therefore compatible with cell biochemistry, as clearly shown here.

We envision EP to be generally compatible with many macromolecular interaction approaches, from mass spectrometry to immunoprecipitation. If combined with suitable chemical modifications of the probes, for instance by introduction of acutely activatable crosslinking groups, EP may become a method of choice for the identification of elusive binding partners (for instance, low affinity substrates of enzymes) with high spatial and temporal resolution. Here, we have demonstrated how the introduction of a farnesyl chain in vitro rescued the kinetochore localization of Spindly in cells treated with a farnesyl transferase inhibitor. In the future, this approach may be extended to other lipid modifications, and may allow monitoring the behavior of only one or a few modified proteins at the time. Another obvious advantage of *in vitro* manipulations of macromolecules is that it allows precision labeling with small fluorescent dyes, potentially bypassing limits associated with tagging target proteins with bulky genetically encoded fluorescent proteins. In this context, EP would expand the toolbox of reagents suitable for live-cell spectroscopic applications such as FLIM-FRET microscopy or FRAP analysis. In summary, we envision that EP has the potential to become a method of choice for delivery of synthetic or semi-synthetic proteins into the cellular environment.

## Methods

### Electroporation of living cells

Electroporation (EP) was performed using either the Neon Transfection System Kit (Thermo Fisher) or the Amaxa Nucleofector I system (Lonza). For EPs with Neon Transfection System, cells were harvested by trypsinisation, washed with PBS, and resuspended in Buffer R (Thermo Fisher) to a final volume of 90 µl. Protein samples were diluted 1:1 in buffer R and then a volume of 30 µl of this mix was added to the cell suspension (the EP slurry). Volumes and protein-buffer ratios may be adjusted according to the purpose of the experiment and depending on protein solubility. Final protein concentrations in the respective EP slurry varied between 5 and 200 µM (in a final volume of 110 μL). The EP slurry was loaded into a 100 µl Neon Pipette Tip (Thermo Fisher) and electroporated with 2 consecutives pulses at 1000V and for durations of 35 msec (with the exception of the experiments in U2OS, RPE and HEK293 cells, which were electroporated with 2 × 25 msec pulses at 800V). Following EP, the slurry was added to 50 ml of pre-warmed PBS, sedimented by a 0.5xg centrifugation and trypsinised for 5 to 7 minutes to remove non-internalized extracellular protein. Following one further wash in PBS and centrifugation, the cell pellet was re-suspended in complete imaging medium (without antibiotics) and transferred to a cell-imaging plate (Ibidi). Cells were returned to the incubator and allowed to recover for a minimum of 5 hrs. After recovery, cells were either analyzed by fluorescence microscopy imaging or processed to generate protein extracts to be used for either immunoprecipitation analysis or western blotting.

EP with the Amaxa system was performed according to the reported protocol (21). EP conditions described above gave satisfactory results for all the cell types and proteins tested. Only a small fraction of the tested recombinant proteins proved unsuitable for EP delivery. This was due to formation of major intracellular aggregates or because the protein samples failed to localize correctly within the cell. We advise to initially vary EP parameters, most notably pulse strength and duration, for optimal performance. Moreover, cell-specific and protein-specific optimization of such parameter should be always carried out in order to achieve the best trade-off between cell viability and efficient protein delivery. The following are some of the important factor to consider during optimization of protein delivery: 1) In our work, we invariably used protein samples of great purity and homogeneity (high monodispersity and solubility). Although we did not systematically analyze the relationship between protein homogeneity and efficiency of EP, we suspect that protein sample quality is an important factor. 2) Centrifugation of protein sample prior to EP is strongly recommended. 3) During the recovery phase, we recommend avoiding the use of selection antibiotics in the media as they increase cell mortality. 4) Once resuspended in EP buffer, cells should be electroporated and washed as fast as possible to reduce cytotoxicity. 5) If required, including additional trypsinization and washing steps following EP may improve the removal of persistent extracellular protein aggregates generated during EP.

### Production of recombinant proteins

All recombinant proteins used in this study were of human origin. For expression and purification of recombinant proteins, synthetic codon-optimized DNA encoding human *MAD2, BUBR1, SPINDLY*, *NDC80C* and *MIS12C* subunits were used. All proteins used in this study were full length, except mTurquoise2-BUBR1^1-571^ (which we refer to as CFP-BUBR1^1-571^) and mTurquoise2-BUBR1^1-571(Δ432–484)^ (which we refer to as CFP-BUBR1^1-571-ΔHelix^). Fluorescent MAD2^TAMRA^, CFP-BUBR1^1-571^, mCherry/GFP-Spindly constructs, NDC80C^WT^-GFP complex and EGFP-K-RAS D30C Tf3 were expressed, purified, and labelled as previously described (1, 32, 37, 57). Recombinant NDC80C^9A^-GFP complex was generated by fusing a C-terminal GFP and a His6-tag to SPC25. Construct for insect cell expression exploited the MultiBac baculovirus expression system (62). A Bacmid was then produced from EMBacY cells and subsequently used to transfect Sf9 cells and produce baculovirus. Baculovirus was amplified through three rounds of amplification and used to infect Tnao38 cells. Cells infected with virus were cultured for 72 h before harvesting. Cell pellets were resuspended in buffer A (50 mM HEPES pH 8, 200 mM NaCl, 20 mM Imidazole, 5% glycerol, 2 mM TCEP) supplemented with protease-inhibitor mix from Serva and 0.2% Triton, lysed by sonication and cleared by centrifugation. NDC80C^9A^-GFP was then eluted in buffer B (50 mM HEPES pH 8, 200 mM NaCl, 300 mM Imidazole, 5% glycerol, 2 mM TCEP) and the eluate was diluted 6 times in volume using ion exchange buffer C (50 mM HEPES pH 8, 25 mM NaCl, 5% glycerol, 2 mM TCEP, 1 mM EDTA) and applied to a 6 mL Resource Q anion-exchange column pre-equilibrated in the same buffer. Elution of bound protein was achieved by a linear gradient (25-400 mM NaCl in 25 column volumes). Relevant fractions were concentrated in 10 kDa molecular mass cut-off Amicon concentrators and applied to a Superose 6 16/70 column equilibrated in size-exclusion chromatography buffer D (50 mM HEPES pH 8, 250 mM NaCl, 5% glycerol, 2 mM TCEP). Peak fractions containing the NDC80C^9A^-GFP complex were collected and further concentrated in a 10 kDa cut-off Amicon concentrator before being flash frozen in liquid N2 and stored at −80°C. Recombinant CFP-MIS12 complex was generated by fusing N-terminal CFP to PMF1. Baculoviral transfer vectors encoding CFP-Mis12C expression were generated using biGBac platform (63). Baculoviruses were generated in Sf9 cells, and expression of CFP-tagged MIS12C was carried out in Tnao38 cells for 72h-96h, at 27°C. CFP-Mis12C was purified by a three-step protocol, as described previously for the non-fluorescent version (39): i) affinity purification of filtered supernatant with 5 ml HisTrap FF column (GE Healthcare) and step elution with 300 mM imidazole; ii) anion-exchange of the dialyzed eluate with 6 ml Resource Q column, elution with linear NaCl gradient; and iii) final polishing step via size-exclusion on a Superdex 200 10/300 column (GE Healthcare) equilibrated in 20 mM Tris-HCl (pH 8.0), 0.15 M NaCl, and 1 mM TCEP. Relevant fractions were pooled, concentrated, flash-frozen in liquid nitrogen and stored at −80°C. Recombinant CFP-BUBR1^1-571-ΔHelix^ was expressed in Tnao38 cells that were then lysed in buffer A (25 mM HEPES (pH 7.5), 300 mM NaCl, 10% glycerol, 2 mM TCEP, 1 mM PMSF). Soluble lysates were passed over a 5-ml Ni-NTA column and, after washing with 20 column volumes buffer A, the proteins were eluted by adding 300 mM imidazole to buffer A. Proteins were subsequently gel-filtered on a Superdex S200 16/60 column equilibrated against buffer B (10 mM HEPES (pH 7.5), 150 mM NaCl, 5% glycerol, 2 mM TCEP). Fractions containing purified protein were concentrated, flash-frozen and stored at −80 °C. Recombinant N-terminally acetylated human α-SYNUCLEIN (αSYN) was purified from *E. coli* as reported (21). In vitro pre-farnesylation of mCherry-Spindly (30 µM) was achieved by incubation with recombinant Farnesyltransferase (10 μM) and Farnesylpyrophosphate (90 µM) for 5-6 hours at 20°C, followed by gel filtration (S200) purification to remove the Farnesyltransferase.

### Cell culture, siRNA transfection, immunoprecipitation, immunoblotting and analysis of intracellular protein levels

The following cell lines were cultured in DMEM (PAN Biotech) supplemented with 10 % FBS (Clontech), penicillin, streptomycin (GIBCO) and 2 mM L-glutamine (PAN Biotech): HeLa, mCherry-H2B HeLa, U2OS, MDCK, HEK293, and RPE. Human A2780, SK-N-SH and SH-SY5Y cells were grown in RPMI 1640, DMEM-Ham’s F-12 and DMEM respectively. Cells were grown at 37 °C in the presence of 5% CO_2_. All experiments requiring live imaging were performed in complemented CO_2_-independent medium (GIBCO) at 37 °C. Cells used in this study are regularly checked for mycoplasma contamination. Cell lines were not authenticated. Unless differently indicated, the microtubule-depolimerising drug nocodazole was used at 3.3 μM (Sigma). Endogenous farnseyltransferase inhibition was achieved at 10 μM of FTI-277 (Sigma). Cellular RAS activity was stimulated with 50ng/ml EGF (Sigma). Were indicated, the DNA dye SiR-Hoechst-647 Dye (Spirochrome) at a concentration of 0.5 µM was added to the medium 1 hour before live imaging. Depletion of endogenous MIS12C was achieved by RNAiMax (Invitrogen) transfection of 3 combined siRNA duplexes used at 10nM each for 48 hrs (RNA oligos sequence for Dsn1 is GUCUAUCAGUGUCGAUUU; for Nsl1 is CAUGAGCUCUUUCUGUUUA; Sigma-Aldrich) (RNA oligos sequence for MIS12 is for GACGUUGACUUUCUUUGAU; GE Healthcare Dharmacon). To generate mitotic populations for immunoprecipitation experiments after EP, cells were treated with nocodazole for 16 hours. Mitotic cells were then harvested by shake off and resuspended in lysis buffer [150 mM KCl, 75 mM Hepes, pH 7.5, 1.5 mM EGTA, 1.5 mM MgCl_2_, 10 % glycerol, and 0.075 % NP-40 supplemented with protease inhibitor cocktail (Serva) and PhosSTOP phosphatase inhibitors (Roche)]. A total of 4 mgs of protein extract per sample was then incubated with GFP-Traps beads (ChromoTek; 3 μl/mg of extract) for 3 hours at 4 °C. Immunoprecipitates were washed with lysis buffer and resuspended in sample buffer, boiled and analyzed by SDS-PAGE and Western blotting using 4-12 % gradient gels (NuPAGE). The following antibodies were used for the western blot analysis in this study: anti-Bub1 (rabbit polyclonal; Abcam; 1:5000), anti-Hec1 (human Ndc80; mouse clone 9G3.23; Gene- Tex, Inc.; 1:250), anti-Mis12 (in house made mouse monoclonal antibody; clone QA21; 1:1000), anti-GFP (in house made rabbit polyclonal antibody; 1:1,000-4,000) anti-Vinculin (mouse monoclonal; clone hVIN-1; Sigma-Aldrich; 1:10000), anti PMF1/NNF1 (in house made mouse affinity purified monoclonal; clone RH25-1-54, 1:1000) and anti-Tubulin (mouse monoclonal, Sigma-Aldrich; 1:10000). Quantification of protein levels from western blots was performed with the following formula: [(LPoIsiRNA-PoIBgr)/(LVincsiRNA-VincBgr)]/[(LPoICtrl-PoIBgr)/(LVincCtrl-VincBgr)]. LPoIsiRNA= levels of the protein of interest for the siRNA lane; PoIBgr= background signal for the protein of interest; LVincsiRNA= levels of Vinculin for the siRNA lane; VincBgr= background signal for Vinculin; LPoICtrl= levels of the protein of interest for the Control lane; PoIBgr= background signal for the protein of interest; LVincCtrl= levels of Vinculin for the Control lane; VincBgr= background signal for Vinculin. Fluorimetric analysis was performed using Greiner flat-bottom plates and a Clariostar microplate reader (monochromator excitation at 587+/− 10 nm, emission 610 +/− 10 nM). Fluorescence intensity from protein extracts derived from a known number of electroporated cells were measured and plotted against a calibration curve generated with defined concentrations of recombinant mCherry. Bar graphs show average intracellular concentrations and SD for two independent experiments in which every concentration was analysed in duplicate.

### α-SYNUCLEIN detection and flow cytometry

For immunofluorescence imaging of delivered αSYN in fixed samples (electroporation concentration 400 µM), cells were recovered for 5 hrs in the incubator on poly-L-lysine-coated 25 mm cover slips. Cells were quickly washed 3 × with complete medium and treated briefly with diluted trypsin/EDTA (0.01 %/ 0.004 %, 40 s, room temperature) to remove non-internalized αSYN, then fixed in PBS containing 4 % (w/v) PFA for 15 min and permeabilized with 0.1 % (v/v) Triton-X in PBS for 3 min. After washing 3 × 10 min with PBS, samples were blocked with 0.13 % (v/v) cold fish skin gelatin (Sigma) in PBS for 1 h. Cells were incubated for 2 hrs with anti-αSYN ab52168 (Abcam, 1:100 dilution) in blocking buffer. After washing 3 × 10 min with PBS, specimens were incubated with anti-mouse IgG Atto647, (Sigma, 1:1000 dilution) and fluorescein isothiocyanate (FITC)-labeled phalloidin (Millipore, 2 mg/mL) for 1 h in blocking buffer. Slides were washed 3 × 10 min with PBS and nuclei stained with 1 µg/mL 4’,6-diamidino-2-phenylindole (DAPI, Invitrogen) in PBS for 15 min. After washing once in PBS, samples were mounted with Immu-Mount (Thermo Scientific). For αSYN detection by western blotting, cell lysates were separated on commercial 4-18 % gradient SDS-PAGE (BioRad), transferred onto PVDF membranes and fixed with 4 % (w/v) paraformaldehyde (PFA) in PBS for 1 h. Membranes were washed 2 × 5 min with PBS and 2 × 5 min with TBS (25 mM Tris, 136.9 mM NaCl, 2.7 mM KCl, pH 7.4). After blocking for 1 h in 5 % (w/v) milk in TBST (0.1 % (v/v) Tween-20 in TBS), membranes were probed with anti-αSYN sc69977 (Santa Cruz, 1:100 dilution) and anti-Actin IgM (Merck Millipore, JLA20, 1:5,000 dilution). Secondary antibodies were HRP-conjugated anti-mouse or anti-rabbit (Sigma, 1:10,000 dilutions). Membranes were developed using SuperSignal West Pico or Femto chemiluminescent substrates (Thermo Scientific). Luminescence signals were detected on a BioRad Molecular Imager and quantified with ImageLab (BioRad). Flow-cytometry and quantification (fluorescence intensity median analysis) of αSYN containing cells were performed with lysine-to-Alex488 fluorophore-coupled recombinant protein as described (21). In brief, N-hydroxysuccinimide (NHS) ester-activated Atto488 fluorescent dye (Sigma Aldrich) was coupled to αSYN lysine side chain-amines in bicarbonate buffer at pH 8.3 according to the manufacturer’s instructions. Covalently modified αSYN was separated from non-reacted dye on a Sephadex G-25 column (Amersham, GE) and concentrated with 6 kDa MW cut-off spin columns (Millipore). The final concentration of the fluorescently labeled protein was measured by UV-VIS spectrophotometry at 280 nm with E: 5690 M^−1^cm^−1^. A correction factor of 0.1 was subtracted to compensate for the intrinsic absorbance of the fluorophore. Cells containing Atto488-tagged protein (530 nm) were detected by flow cytometry on a FACSCalibur (BD Biosciences, >10,000 recorded events for each sample). Median αSYN fluorescence from all events was determined with FlowJo 8.8.6 for each sample.

### Microscopy, immunofluorescence detection, live-cell Imaging, FRAP analysis, FRET-FLIM imaging and chemical inhibition

For this study, the following microscopes were used. 1) A customized 3i Marianas system (Intelligent Imaging Innovations) equipped with a Axio Observer Z1 microscope chassis (Zeiss), a CSU-X1 confocal scanner unit (Yokogawa Electric Corporation), Plan-Apochromat 63×/1.4 NA objectives (Zeiss), an Orca Flash 4.0 sCMOS Camera (Hamamatsu), and a FRAP Vector™ High Speed Point Scanner. Images were acquired as *Z* sections (using Slidebook Software 6 from Intelligent Imaging Innovations) and converted into maximal-intensity-projection TIFF files for illustrative purposes. 2) A Deltavision Elite System (GE Healthcare) equipped with an IX-71 inverted microscope (Olympus), a UPlanFLN 40×/1.3 NA objective (Olympus) and a pco.edge sCMOS camera (PCO-TECH Inc.). Images were acquired as *Z* sections (using the softWoRx software from Deltavision) and converted into maximal-intensity-projection TIFF files for illustrative purposes. 3) An Olympus FV1000 FlouView laser scanning confocal microscope counted in a single-photon counting avalanche photodiode (PDM Series, MPD; PicoQuant) and timed by using a time-correlated single-photon counting module (PicoHarp 300; PicoQuant) after being spectrally filtered using a narrow-band emission filter (HQ 525/15; Chroma). The microscope was also equipped with an Eppendorf Transjector 5246 and an Eppendorf Micromanipulator 5171. 4) For αSYN experiments, confocal images were taken at 40 × magnifications and using excitation wavelengths of 405, 488, and 633 nm on a Zeiss LSM 510 META laser-scanning microscope. For immunofluorescence analysis, cells (HeLa) were grown on coverslips pre-coated with 15 µg/ml poly-d-lysine (Millipore) and then fixed with 4% paraformaldehyde in PBS, followed by permeabilization with PBS/PHEM-Tween 0.3%. The following antibodies were used for immunostaining: anti-CREST/anti-centromere antibodies (human, Antibodies Inc.; 1:100) anti-Tubulin (mouse monoclonal, Sigma; 1:5000). DNA was stained with 0.5 µg/ml DAPI (Serva) and coverslips mounted with Mowiol mounting medium (Calbiochem). The individual electroporation of HeLa cell with either the CFP-BUBR1^1-571^ constructs or MAD2^TAMRA^ was performed using an EP concentration of respectively 10 µM and 5 µM. After electroporation cells were allowed to recover overnight and then treated with nocodazole for imaging on the Mariana system. To determine CFP kinetochore levels for Figure 1D, between 263and 372 KTs per condition were scored. Simultaneous electroporation of 10 µM CFP-BUBR1^1-571^ and 5 µM MAD2^TAMRA^ was performed on HeLa cells and imaged live on a Deltavision system. To measure mitotic duration in cells depleted for endogenous MIS12C and electroporated with recombinant CFP-MIS12C at 8 µM, H2B-mCherry HeLa cells were transfected with siRNA-MIS12C oligos, twice in 24 hours, and then electroporated 6 hours after the second transfection. Following EP, cells recovered overnight before being processed for either western blotting analysis or imaged lived on the Deltavision system. Mitotic duration, with or without 330 nM nocodazole, was assessed by DNA morphology. Cells entering anaphase with their DNA not completely aligned in the metaphase plate were scored as erroneous segregations. Between 53 and 149 cells per condition were scored. Live-cell time-lapse imaging of HeLa cells electroporated with NDC80C^WT^-GFP at 5 µM was performed on an asynchronously growing population. Cells were then left to recover overnight before being treated with SiR-Hoechst-647 Dye for one hour before being imaged live on a Deltavision system. To determine mitotic duration, between 60 and 70 cells per condition were scored. For the error correction assay, following electroporation HeLa cells were synchronized in G2 with 9 µM RO-3306 (Millipore) for 16 hours and then released in the presence of 5 µM STLC (Sigma-Aldrich) for 3 hours. Following STLC wash-out, cells were grown for 150 minutes in media containing 5 µM MG132 (Calbiochem), then fixed, prepared for immunofluorescence analysis and then scored for the presence of uncongressed chromosomes on the Mariana microscope. Cells with one or more chromosomes not perfectly aligned at the metaphase plate were scored as uncongressed. Results are representing the average ± SD of two replicated experiments. In total, between 140 and 191 cells were scored for each condition. Electroporation of NDC80C^WT^-GFP into RPE1, U2OS or HEK293 cells, was performed on asynchronously growing cultures. Following electroporation cells were left recover overnight and then incubated in nocodazole and SiR-Hoechst-647 Dye for 4 hours, and then imaged live on the Mariana microscope. RPE1 and HEK293 cells displayed a higher cell mortality than other cell lines, thus, for these lines, we recommend a protein-specific optimisation of EP parameters in order to achieve proper cell viability. For Spindly experiments, asynchronously growing HeLa cells were treated with farnesyltransferase inhibitor FTI for 24 hours before being electroporated with fluorescent Spindly at 10 µM. Following EP, cells were left to recover for 24 hours in FTI-277, of which the last 6 hours were in the presence of nocodazole. Immunofluorescence analysis was performed on the Mariana system. To determine kinetochore levels for fluorescent SPINDLY, between 60 and 220 KTs per condition were scored. For EGFP-KRAS D30C-TF3 experiments, microinjections and electroporations for FLIM-FRET imaging were performed on MDCK cells that were serum-starved overnight in MEM Eagle medium (Sigma) supplemented with 0.5% FBS. Following starvation, cells were either microinjected or electroporated with EGFP-KRAS D30C-TF3 at a concentration of 6–10 mg/mL and then left recover for 6 hours before live-cell imaging by FLIM-FRET microscopy. Microinjections and FLIM-FRET imaging were performed on an Olympus FV1000 as previously described (37). FLIM-FRET analysis of EGFP-KRAS D30C-TF3 lifetime was performed on 12 electroporated cells.

## Acknowledgments

We thank Alex Faesen, Melina Schuh, Dean Clift, and all members of the Selenko and Musacchio laboratories for helpful discussions and comments. We thank Pim Joannes Huis in’t Veld for his support in the generation of the GFP-NDC80C-9A mutant. We thank Philippe Bastiaens for support with the KRAS FLIM-FRET analysis. A.M. gratefully acknowledges funding by the Max Planck Society and the European Research Council (ERC) Advanced Investigator Grant RECEPIANCE (proposal number n° 669686).

## Supplemental Figure Legends

**Figure S1.**
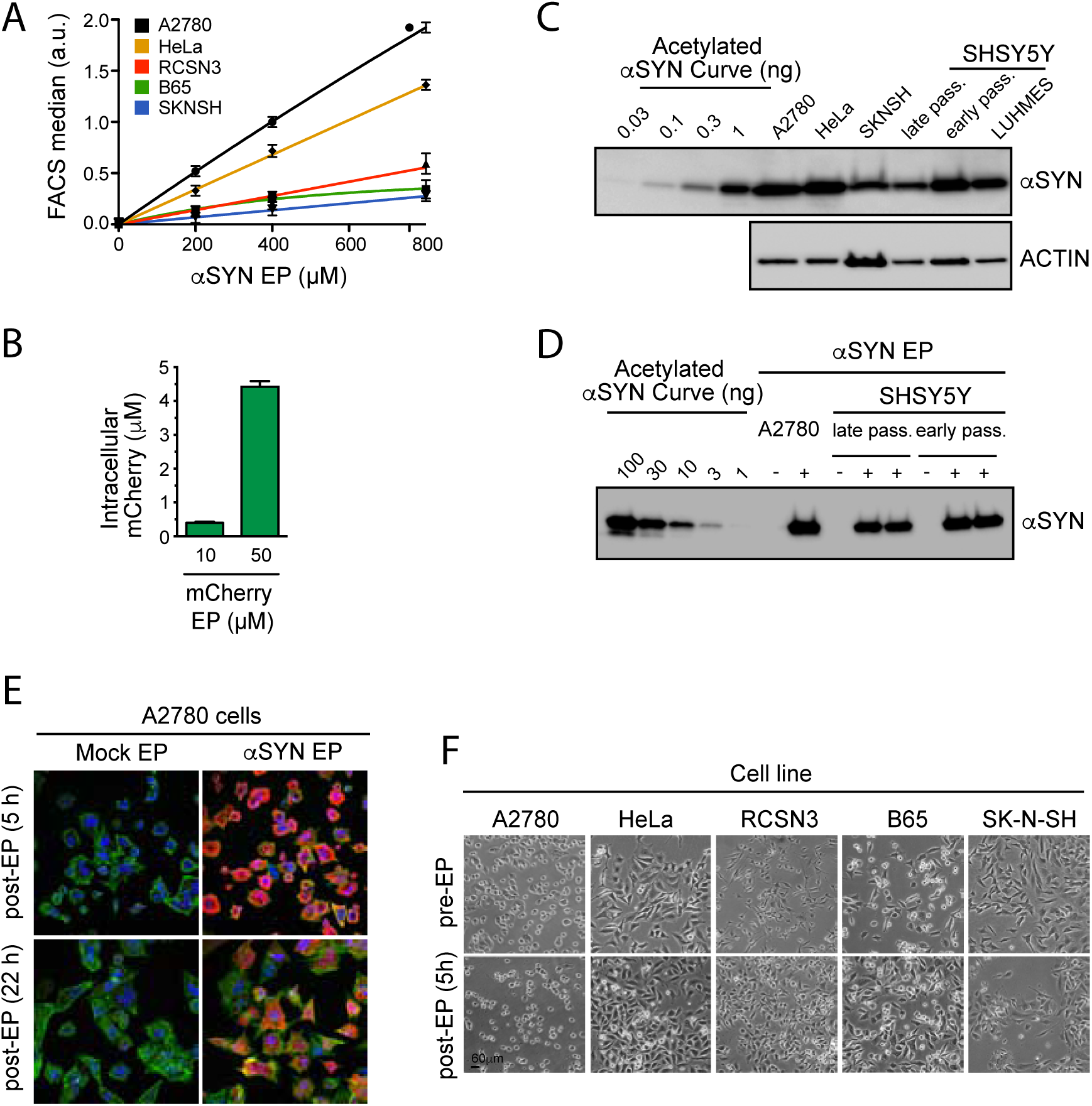
Analysis of the levels of protein uptake and cytotoxicity in electroporated cells. **A**) Quantification of EP-mediated protein uptake by flow cytometry across different cell lines. Median fluorescence intensities (MFI) determined by flow cytometry of EP-processed cells harboring Atto488-labelled α-synuclein (αSYN) plotted against αSYN input concentrations in the respective EP mixtures. Linear uptake efficiencies are evident for all cell lines despite cell-type specific differences in general uptake properties. MFI values were corrected for average cell sizes and corresponding volumes. Lines correspond to linear fits connecting individual measurement points. Error bars are SDM from 4 independent samples. **B**) Quantification of EP-mediated protein uptake by fluorimetric analysis. Extracts from HeLa cells electroporated with mCherry and analyzed with a fluorimeter show that uptake is proportional to the input protein concentration. Fluorescence intensity of protein extracts generated from a known number of cells that were electroporated with different amounts of recombinant mCherry were quantified with a fluorimeter and plotted against an input titration curve. The values obtained were subtracted for the intensity values measured from a mock EP sample. Graph represents mean values ± SD. **C**) Western blotting analysis of whole cell lysates to detect endogenous αSYN levels in A2780, HeLa, SK-N-SH, early- and late-passage SH-SY5Y and undifferentiated LUHMES cells. Defined concentrations of recombinant N-terminally acetylated αSYN serve as input control. Average intracellular concentrations are in the range of 0.05 to 0.5 µM, calculated based on experimentally determined cell volumes (21). **D**) By comparison to the endogenous levels determined in panel C, cells electroporated with 400 µM αSYN in the EP slurry harbor ~100-fold higher levels of the recombinant protein (see input concentrations for reference). **E**) Electroporation of αSYN in A2780 cells produces low levels of cell damage. Panels show anti-αSYN immunofluorescence detection in A2780 cells at different time-points following batch EP and recovery. DNA is shown in blue (DAPI), actin in green (phalloidin) and αSYN in red. Note that most cells harbor αSYN (high penetrance) and that intracellular protein levels are similar between cells (uniform delivery efficiency). 22 hours after EP, adherent cells regain their physiological cell morphologies, an indication that EP caused low cytotoxicity. Mock EP was carried out without recombinant αSYN in the EP mixture. **F**) EP does not have major cytotoxic effects across different cell lines. Bright field analysis of neuronal (RCSN-3, B65, SK-N-SH) and non-neuronal (A2780, HeLa) cell lines electroporated with α-synuclein (αSYN) to inspect cell morphology and viability.

**Figure S2.**
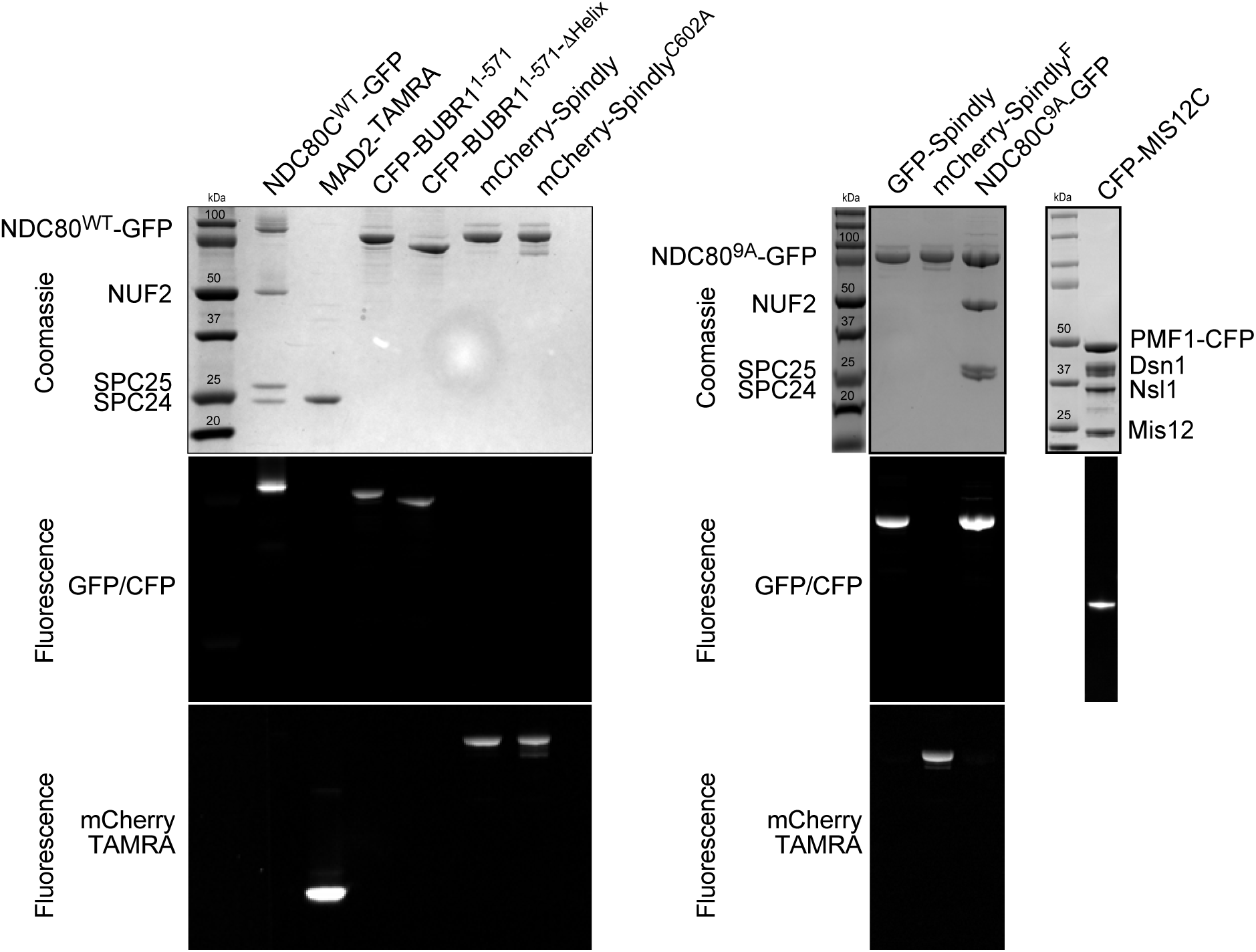
Recombinant proteins used for this study. Protein samples employed in this study were run on an acrylamide gel and imaged by coomassie staining or for the indicated fluorophore. Note: the gel representing fluorescent CFP-MIS12 was run with a sample that was not boiled (to preserve fluorescence) and this resulted in a faster migration.

**Figure S3.**
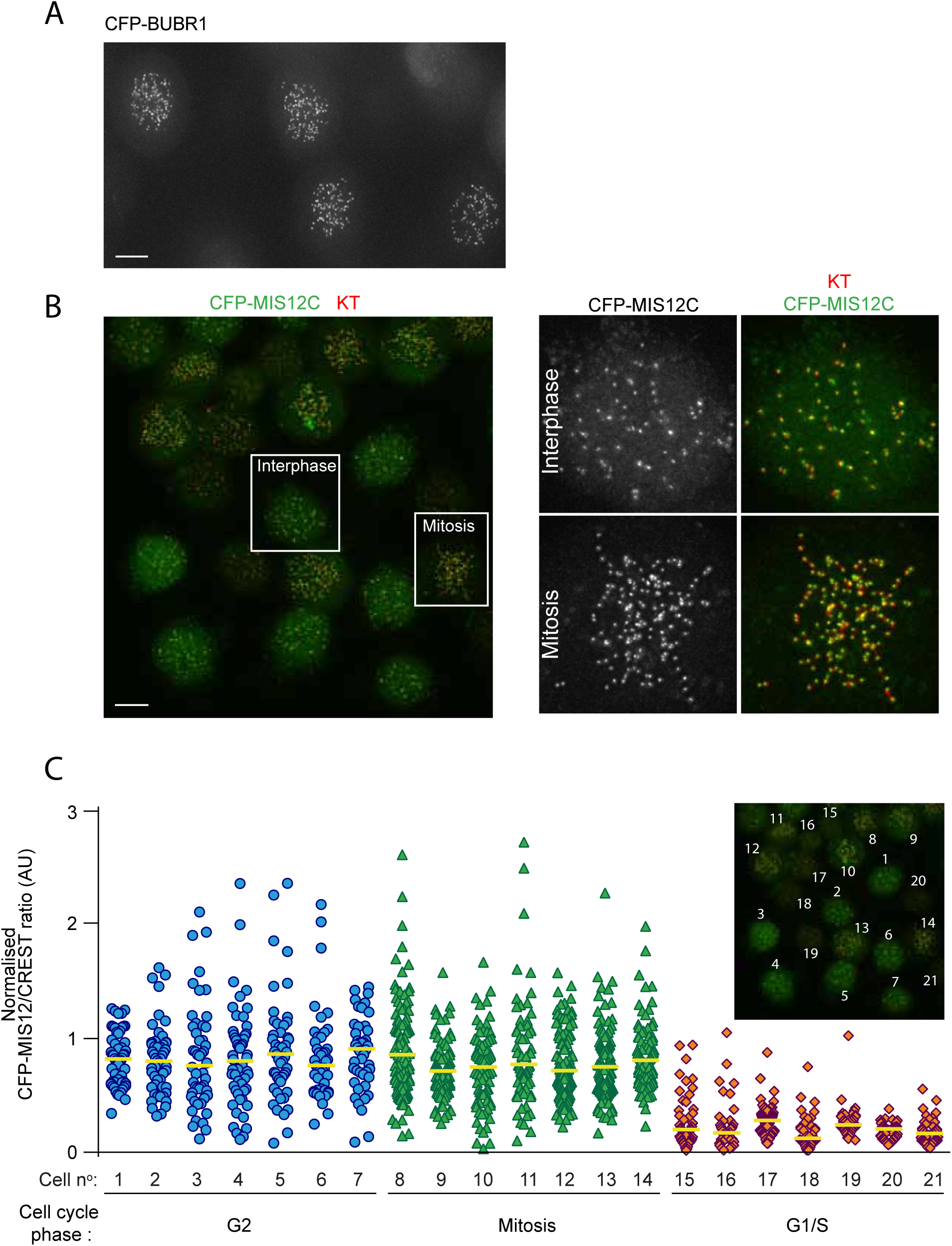
EP is highly efficient and produces uniform protein levels between different recipient cells. A representative imaging field shows HeLa cells electroporated with either CFP-BUBR1 **(A)** or CFP-MIS12C **(B)** and, after recovery, treated with nocodazole for 4 hours. Insets in B represent magnification of the indicated cells. KTs were stained with the marker CREST. Scale bar = 5 μm. **(C)** Quantification of KT levels for electroporated CFP-MIS12C from cells in panel B. Individual cell numbers are indicate in the small inset. Cell phases were assigned based on cell morphology and size. Each symbol represents a single KT. Yellow lines indicate median.

**Figure S4.**
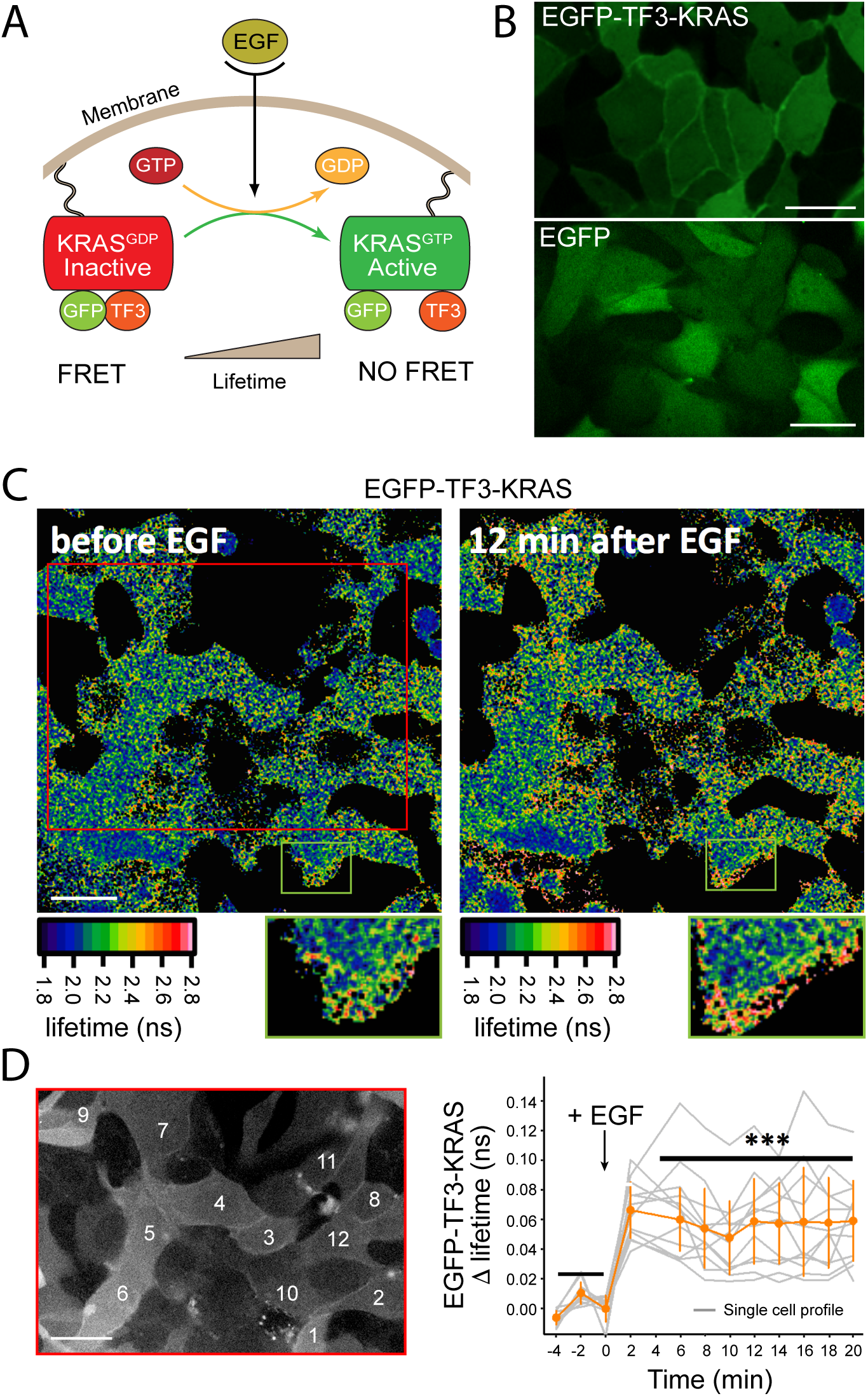
Live-cell FLIM-FRET on EP-delivered proteins. **A**) Schematic representation of the activation of the membrane-bound EGFP-TF3-KRAS FRET-sensor. Treatment with EGF results in a conformational change which results in a loss of FRET between the fluorescent moieties of the sensor and an increase in the donor fluorescence lifetime measured by FLIM-FRET live microscopy. **B)** Membrane localization of EP-delivered EGFP-TF3-KRAS in serum-starved MDCK interphase cells. Scale bar = 5 μm. **C)** Analysis of spatiotemporal dynamics of electroporated EGFP-TF3-KRAS signaling by live-cell FLIM-FRET imaging. Upon EGF stimulation, the conformational switch in intracellular EGFP-TF3-KRAS results in loss of FRET signal and a subsequent increase in the lifetime of the GFP-donor. Lifetime values for each pixel are shown color-coded. Scale bar = 5 μm. **D)** Activity of electroporated EGFP-TF3-KRAS is qualitatively and quantitatively identical to the one obtained in EGFP-TF3-KRAS microinjection experiments (37). Graph shows single cell readouts of fluorescence lifetime profiles. Grey traces represent individual measurements while orange line represents the mean. Error bars are SDM. Scale bar = 5 μm.

**Supplementary Movie-1-4:** Live imaging movies of cells shown in Figure 2A (Ctrl+CFP, RNA+CFP example 1, RNA+CFP example 2, and RNA+CFP-MIS12C respectively). Images show GFP-signal in green and DNA signal in red.

**Supplementary Movie 5** and **6:** Live-imaging of asynchronous HeLa cells electroporated with either recombinant GFP (6) or NDC80C^WT^-GFP (7). Fluorescence time-lapse images show GFP signal (left panel), DNA signal (central panel) and a merge (right panel).

**Supplementary Movie 7-9:** Live imaging movies of cells shown in Figure 3A-B (Mock, GFP and NDC80C^WT^-GFP respectively). Images show GFP-signal in green and DNA signal in red.

